# Rational design of a protein-protein interaction inhibitor that activates Protein Tyrosine Phosphatase 1B

**DOI:** 10.64898/2026.03.19.712938

**Authors:** Avinash D. Londhe, Sophie Rizzo, Syed M. Rizvi, Alexandre Bergeron, R. Sudheer Sagabala, Nilesh K. Banavali, Damien Thévenin, Benoit Boivin

**Affiliations:** Department of Nanoscale Science and Engineering and Department of Biological Sciences, University at Albany, Albany, NY 12222; Department of Chemistry, Lehigh University, Bethlehem, PA 18015; Montreal Heart Institute Research Center, Montréal, QC H1T 1C8, Canada; Division of Translational Medicine, Wadsworth Center, New York State Department of Health, Albany, NY 12237

**Keywords:** Protein Tyrosine Phosphatases, PTP1B, cell signaling, redox, protein-protein interaction inhibitor

## Abstract

Reversible inactivation of protein tyrosine phosphatases by reactive oxygen species (ROS) is essential to the phosphorylation of growth factor receptors. An important outcome of the inactivation of protein tyrosine phosphatase 1B (PTP1B) by ROS involves the conformational change of its phosphotyrosine binding loop which adopts a solvent exposed position in its oxidized form. We previously demonstrated that 14-3-3ζ binds to the phosphotyrosine binding loop of the oxidized form of PTP1B. Using a rational approach, we developed a unique protein-protein interaction (PPI) inhibitor peptide derived from the phosphotyrosine binding loop of PTP1B designed to disrupt the interaction between PTP1B and the 14-3-3ζ-complex. Exploiting this cell-permeable peptide, we showed decreased association between PTP1B and the 14-3-3ζ-complex in cells treated with epidermal growth factor (EGF). We also demonstrated that preventing the association of this 14-3-3ζ-complex to PTP1B deterred oxidation and inactivation of PTP1B following EGF receptor (EGFR) activation and generation of ROS. Treating cells with our PPI inhibitor decreased EGFR phosphorylation on PTP1B-specific sites. Furthermore, treating EGFR-driven epidermal cancer cells with our PPI inhibitor also significantly inhibited colony formation and cell viability, consitent with increased activation of PTP1B. These data highlight the ability of PTP1B to downregulate critical signaling pathways in cancer when activated using peptide drugs such as our protein-protein interaction inhibitor. We anticipate that preventing or destabilizing the reversible oxidation of other members of the protein tyrosine phosphatase superfamily using PPI inhibitors may offer a foundation for a broad therapeutic approach to rectify dysregulated signaling pathways *in vivo*.

**Significance Statement:** Limited understanding of redox mechanisms regulating PTP catalytic activity is a major knowledge gap that has hampered our efforts to develop activation strategies. In its reversibly oxidized and inactivated form, conformational changes of PTP1B influence its association with regulatory proteins. We demonstrate that designing a cell-permeable peptide based on a loop of PTP1B that becomes exposed during oxidation can block its interaction with the 14-3-3ζ-multiprotein complex and activate the phosphatase. Moreover, activating PTP1B using our protein-protein interaction inhibitor peptide decreases the phosphorylation of its substrate EGFR and decreases the effectiveness of cancer cells to form colonies. This study provides important insights into the therapeutic potential of protein-protein interaction inhibitors that regulate the redox cycle of PTPs to reestablish physiological signaling.

## Introduction

The initial burst of tyrosine phosphorylation that occurs following growth factor-induced activation of tyrosine kinase receptors (RTKs) is sustained by the activation of specific oxidases (1, 2). When reduced nicotinamide adenine dinucleotide phosphate oxidases (NOXs) are activated by RTKs, they commonly generate a controlled amount of superoxide and its dismutated product, hydrogen peroxide (H_2_O_2_), which in turn transiently inhibit members of the protein tyrosine phosphatase (PTP) superfamily (3–7). Interestingly, previous studies have also shown that prolonged inactivation of specific PTPs by these cellular oxidants directly contributes to increased tyrosine-dependent signaling in several diseases (7–11). Although these observations point to potential therapeutic applications of PTP activation, most drug discovery efforts to restore tyrosine phosphorylation-dependent signaling to physiological levels have focused on protein tyrosine kinases, as reflected by the number of FDA-approved tyrosine kinase inhibitors now exceeding 70 (12).

While the ∼100 members of the PTPome present a wide structural diversity, they all share a conserved HC(X)₅R signature motif positioned on the PTP loop at the bottom of a surface-accessible active-site cleft (13). The architecture of this cleft confers an unusually low pKa to the conserved cysteine residue within this motif, enabling it to maintain a deprotonated state, as a thiolate ion, at physiological pH (14). The reactive, nucleophilic, nature of the catalytic cysteine is central to the ability of PTPs to attack the phosphorus on their substrate in the dephosphorylation process. However, it also makes it prone to react with cellular oxidants, with the oxidation of the catalytic cysteine acting as a transient redox switch to inhibit PTP activity (6).

The structure of protein tyrosine phosphatase 1B (PTP1B), the first PTP to be crystalized in its reversibly oxidized form, revealed several striking conformational changes at the active site of the enzyme (15–17). At the center of these alterations, the crystal revealed that the reversible oxidation of the catalytic cysteine residue led to the formation of a cyclic sulfenamide with the adjacent serine residue, severely distorting the PTP loop. The constrain imposed by the formation of the cyclic sulfenamide weakens several interactions and triggers the PTP loop, usually buried in the active site, to flip out and to become solvent exposed. Another major change that occurs as a consequence of the PTP becoming reversibly oxidized is seen in the phosphotyrosine (pTyr) binding loop that relocates from the edge of the active-site cleft, flipping over to expose a sequence of ten amino acids (Lys^41^-Ser^50^) to the cytosol. With the pTyr loop pushing up approximately 9 Å (18), this conformational change exposes a serine that can be phosphorylated by AKT (19). Our previous work has shown that 14-3-3ζ binds to the newly exposed and phosphorylated pTyr loop in the reversibly oxidized form of PTP1B (PTP1B-OX) to transiently stabilize the phosphatase in its inactivated form (18).

Herein, we took advantage of the conformational changes occurring at the active site of PTP1B-OX to develop a protein-protein interaction (PPI) inhibitor designed to prevent the interaction between PTP1B-OX and 14-3-3ζ. In testing this hypothesis, we demonstrated that preventing the interaction between PTP1B and a 14-3-3ζ-complex could activate the phosphatase. We observed that a cell-permeable peptide, derived from the pTyr loop of PTP1B could activate the phosphatase, cause the dephosphorylation of its substrate EGFR and prevent cancer cells to form colonies, a hallmark of carcinogenesis. These findings delineate a previously unexploited opportunity to modulate tyrosine kinase-dependent signalling through the PTP family and provide a conceptual framework for developing PPIs and small-molecule activators of PTP1B and related PTPs.

## Results

### A Cell-Permeable Peptide Disrupts the Interaction Between PTP1B and 14-3-3ζ

Our previous work shows that the rearrangement of the active site cleft that cause the pTyr loop of PTP1B-OX to become exposed to the cytosol (Fig. S1A, B), leads to its association with 14-3-3ζ and to the stabilization of the reversibly inhibited form of the enzyme (18). Our proteomic data also revealed that although 14-3-3ζ mostly associated with the Ser^50^-phosphorylated form of the pTyr loop peptide (Lys^41^-Ser^50^), that it was also associated with the non-phospho peptide in lysates. To investigate this interaction, peptides comprising Lys^41^ to Ser^50^ of PTP1B, or a similar peptide phosphorylated on Ser^50^ were synthesized and immobilized on beads to be used as bait in cell lysates (Fig. 1A). Our pulldowns show that 14-3-3ζ was associated with the non-phosphorylated pTyr loop peptide when compared to control beads (Fig. 1B), however, this enrichment was greatly increased increased in the phosphoSer^50^ peptide pulldowns. Interestingly, when directly assessing the association between purified 14-3-3ζ and the pTyr loop peptides *in vitro*, the interaction was specific for the phosphoSer^50^ peptide (Fig. 1C). Taken together, this suggests that while the interaction between PTP1B and 14-3-3ζ is specific when the phosphatase is oxidized and phosphorylated on Ser^50^, 14-3-3ζ is also interacting with the pTyr loop as part of a larger 14-3-3ζ complex.

**Fig. 1.**
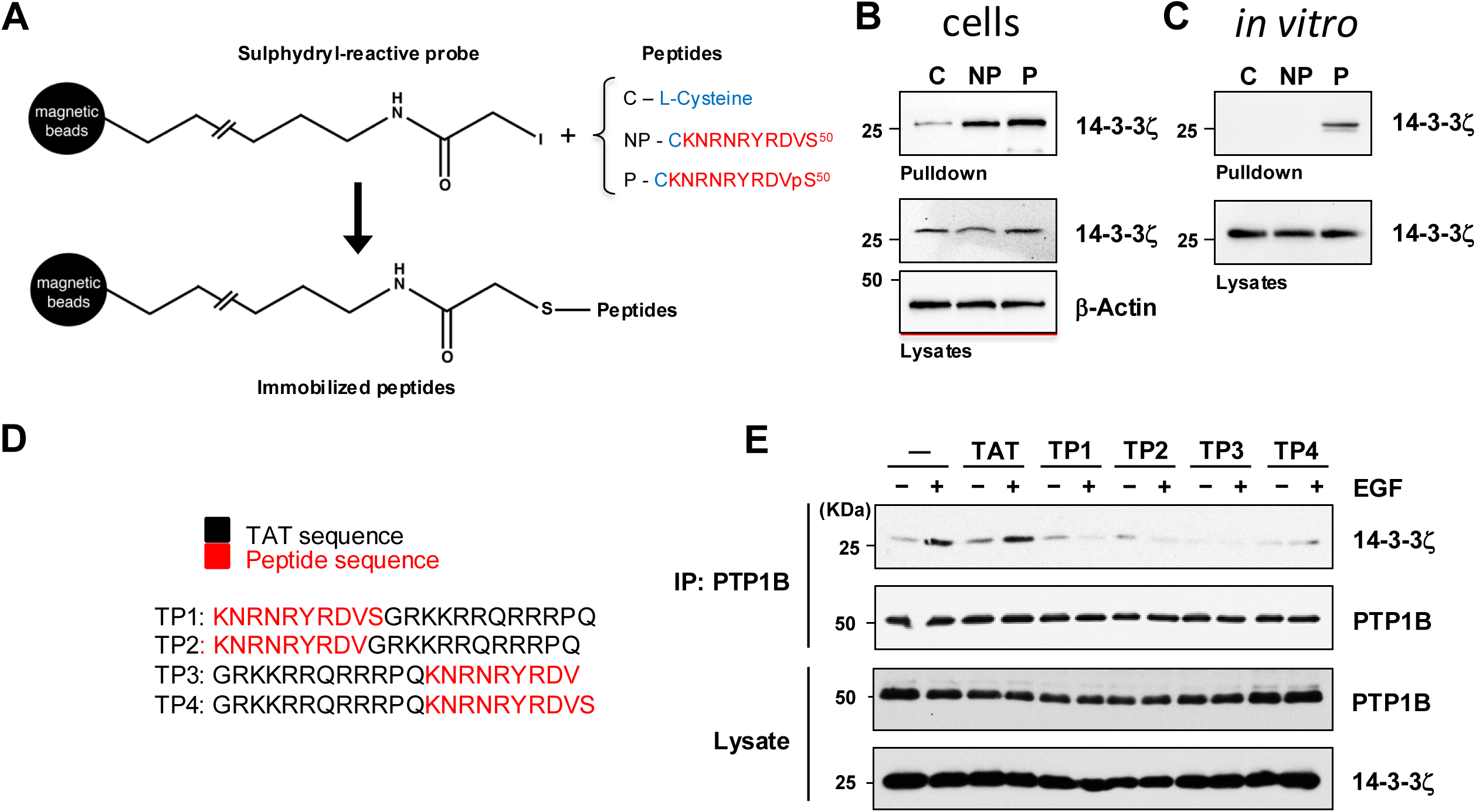
Cell-permeable PTP1B-derived peptides prevent the association between PTP1B and a 14-3-3ζ-protein complex. (*A*) Schematic of the sulphydryl-reactive probe and peptides used to study proteins that interact with the phosphotyrosine binding loop of PTP1B (pTyr loop, Lys^41^-Ser^50^). HEK293T cell lysates (*B*) purified 14-3-3ζ (*C*) were either incubated with magnetic beads linked to L-Cysteine (Cys), the non-phospho pTyr loop-derived peptide (NP) or to a phosphoSer^50^ pTyr loop-derived peptide (P). Pulldowns and lysates were blotted for 14-3-3ζ and β-actin. This experiment was repeated three independent times with representative data shown. (*C*) Design of four cell-permeable <u>T</u>AT-phospho-tyrosine recognition loop <u>p</u>eptides (TP1, 2, 3, 4). (*D*) Lysates from serum-deprived HEK293T cells, stimulated with EGF (100 ng/ml) for 0 or 2 minutes in presence or in absence of TP1, 2, 3 and 4 (15 μM, 60 min) were utilized for PTP1B immunoprecipitations using anti-Flag antibodies. Proteins were separated by SDS-PAGE and probed for 14-3-3ζ using anti-HA antibodies. Lysates were probed for 14-3-3ζ and PTP1B to control for protein expression. This experiment was repeated three independent times with representative data shown.

Based on this finding and the knowledge that the residues of the pTyr loop are only partially conserved between members of the PTP family (13), we designed cell-permeable pTyr loop-derived peptides from the sequence of PTP1B [Fig. 1D, TAT-Peptide (TP) 1, 2, 3 and 4] to destabilize the interaction between PTP1B-OX and the 14-3-3ζ complex. These peptides possess a TAT sequence either in carboxy-(TP1, 2) or amino-terminal (TP3, 4), and encode Ser^50^ (TP1, 4) or not (TP2, 3). We tested whether these peptides could act as PPI inhibitors in HEK293 cells when the association between PTP1B-OX and a 14-3-3ζ-complex is maximal, at two minutes following EGF stimulation (18). While TAT alone showed no effect on the interaction between PTP1B and 14-3-3ζ, all four TAT-Peptides prevented the interaction (Fig. 1D). Hence, peptides derived from the pTyr loop of PTP1B can disrupt the interaction between PTP1B-OX and the 14-3-3ζ-complex whether Ser^50^ is present or not, suggesting that the binding of the complex precedes a specific 14-3-3ζ association with PTP1B-OX.

### Disrupting the Interaction Between PTP1B and the 14-3-3ζ-Complex Activates PTP1B

Although all four TAT-Peptides disrupted the interaction between PTP1B-OX and the 14-3-3ζ-complex, the shared amino-terminal positioning of the TAT moiety –as seen with the sulfhydryl-reactive probe used as bait in Fig. 1– and the presence of Ser⁵⁰ led us to prioritize TAT-Peptide 4 for further investigation. To elucidate the functional role of the PTP1B-OX–14-3-3ζ-complex interaction and to assess whether disrupting this association influences PTP1B activity, we exposed cells to our TP4 disrupting peptide. As expected from results shown in Figure 1D, exposing cells to TP4 prevented the interaction between PTP1B and the 14-3-3ζ-complex following EGFR activation (Fig. 2A). We then assessed the endogenous catalytic activity of PTP1B in lysates from EGF-stimulated cells that were pretreated with TP4 or not, to determine whether disrupting the complex could affect the catalytic activity of PTP1B. PTP1B was immunoprecipitated in oxygen-free conditions to prevent post-lysis oxidation, and the activity of immune-complexes was measured using a pTyr analog, pNPP, as substrate (20). We observed that PTP1B activity transiently decreased to ∼ 30% in lysates from cells exposed to EGF for 2 minutes before recovering at the 10 minutes time point (Fig. 2B). On the other hand, PTP1B inactivation was not observed at 2 minutes in cells that were pre-treated with TP4 prior to EGF stimulation. This suggests that the reversible oxidation of PTP1B is affected by our PPI inhibitor peptide. To assess this possibility, we performed a cysteinyl-labeling assay, a 3-step assay that converts the reversible oxidation of PTPs to a modification by biotin that can be visualized by immunoblotting after a biotin-streptavidin purification step (Fig. 2C) (5, 21). As shown in Figure 2D, biotinylation of PTP1B was detected at 1, 2 and 5 minutes following EGF exposure, however, PTP1B was not biotinylated in cells that were pretreated with TP4. These results were also validated by measuring the presence of active PTP1B using a direct cysteinyl labeling approach (Fig. S2). Hence, breaking the interaction between PTP1B and the 14-3-3ζ-complex revealed some activating properties of our PPI inhibitor by destabilizing or impeding the reversible oxidation of PTP1B.

**Fig. 2.**
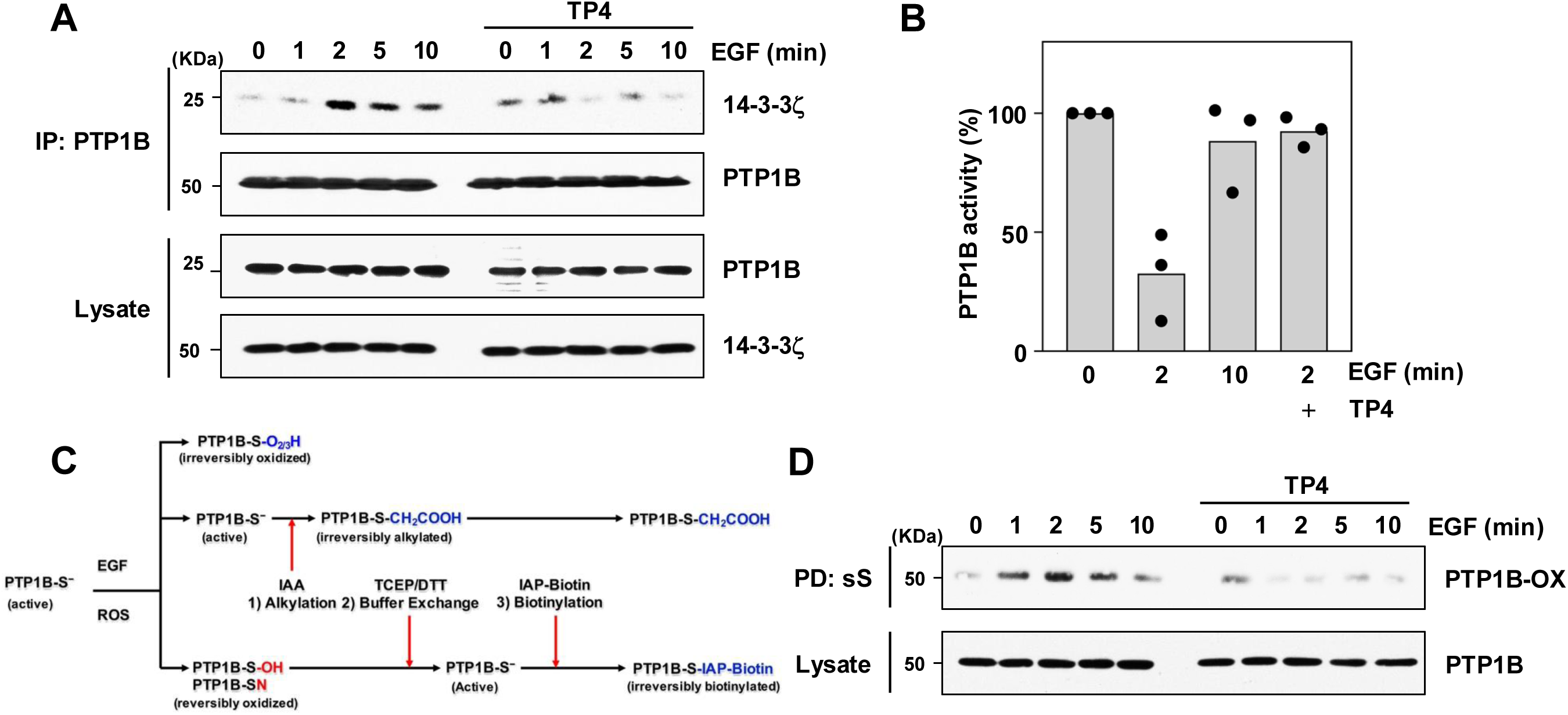
Disrupting the interaction between PTP1B and 14-3-3ζ activates PTP1B. (*A*) HEK293T cells pre-treated or not with TP4 were stimulated with EGF for the indicated times. PTP1B was immunoprecipitated and proteins were separated by SDS-PAGE and probed for 14-3-3ζ. Lysates were probed for 14-3-3ζ and PTP1B to control for protein expression. This experiment was repeated three independent times with representative data shown. (*B*) PTP1B was immunoprecipitated from HEK293T cell lysates in the presence or absence of TP4, stimulated with EGF for the indicated times. Catalytic activity of PTP1B from immune-complexes was measured using pNPP in absence of reducing agent to monitor intracellular PTP1B activity. The average readout value of technical replicates is represented by the dot-plot bar graph. N = 3 independent experiments are shown. (*C*) The cysteinyl-labeling assay is a 3-step assay designed to measure reversible oxidation of PTPs. In the first step of the assay, the pool of PTP1B that remained active following cell stimulation react and is irreversibly alkylated by IAA, whereas all oxidized forms of PTP1B are unreactive and protected from alkylation. Excess IAA is removed in the second step and the pool of reversibly oxidized PTP1B is reduced and reactivated back to their thiolate form using reducing agents (TCEP or DTT). In the third step of the assay, reactivated PTP1B is labeled by a biotinylated thiolate-reactive probe (IAP-Biotin) that allows purification by streptavidin pull-down. (*D*) Reversible-oxidation of PTP1B was measured following TP4 pretreatment using the cysteinyl-labeling assay. HEK293T cells were pre-incubated or not with TP4, stimulated with EGF for the indicated times, and subjected to the cysteinyl-labeling assay. Biotinylated proteins were purified on streptavidin–Sepharose beads, resolved by SDS-PAGE, and blotted for PTP1B. Lysates were probed for PTP1B to control for protein expression and loading. This experiment was repeated three independent times with representative data shown.

### TP4 Prevents Hydrogen Peroxide-Induced PTP1B-14-3-3ζ Complex Formation

Activation of growth factor receptors, such as EGFR, lead to the assembly of NOXs, the generation of superoxide, and its rapid conversion into H_2_O_2_ (22, 23). Since the intracellular oxidation of PTPs occurs primarily via the controlled generation of H_2_O_2_ downstream of activated growth factor receptors (7), we next assessed whether the association between PTP1B and the 14-3-3ζ-complex was specific to EGFR-mediated redox signaling, or whether H_2_O_2_ itself could also trigger this association. As anticipated, in cells exposed to EGF, PTP1B was associated with the 14-3-3ζ-complex at 1, 2 and 5 minutes, with the interaction peaking at 2 minutes (Fig. 3A). However, H_2_O_2_ applied to the extracellular milieu caused a slow but gradual increased interaction between PTP1B and the 14-3-3ζ-complex that culminated at 10 minutes, suggesting that PTP1B-OX is similarly interacting with the 14-3-3ζ-complex following cell exposure to H_2_O_2_. We next pretreated cells with TP4 prior to H_2_O_2_ exposure. Co-immunoprecipitations in these conditions revealed that the interaction between PTP1B and the 14-3-3ζ-complex could be disrupted by our PPI inhibitor in cells exposed to H_2_O_2_ as in EGF-stimulated cells (Fig. 3B). Finally, since H_2_O_2_ signaling is key to PTP1B oxidation and to the assembly of the PTP1B-14-3-3ζ-complex, we tested whether exposing cells to TP4 affected H_2_O_2_ production. Using dichlorodihydrofluorescein diacetate (DCFH-DA), which is converted to a fluorescent derivative (2’,7’-dichlorofluorescein), DCF) in presence of H_2_O_2_, we observed an EGF-induced increase in ROS that was unaffected by pre-incubating cells with TAT alone or TP4 (Fig. 3C). This is consistent with TP4 preventing the interaction between PTP1B and the 14-3-3ζ-complex without altering growth-factor mediated H_2_O_2_ generation.

**Fig. 3.**
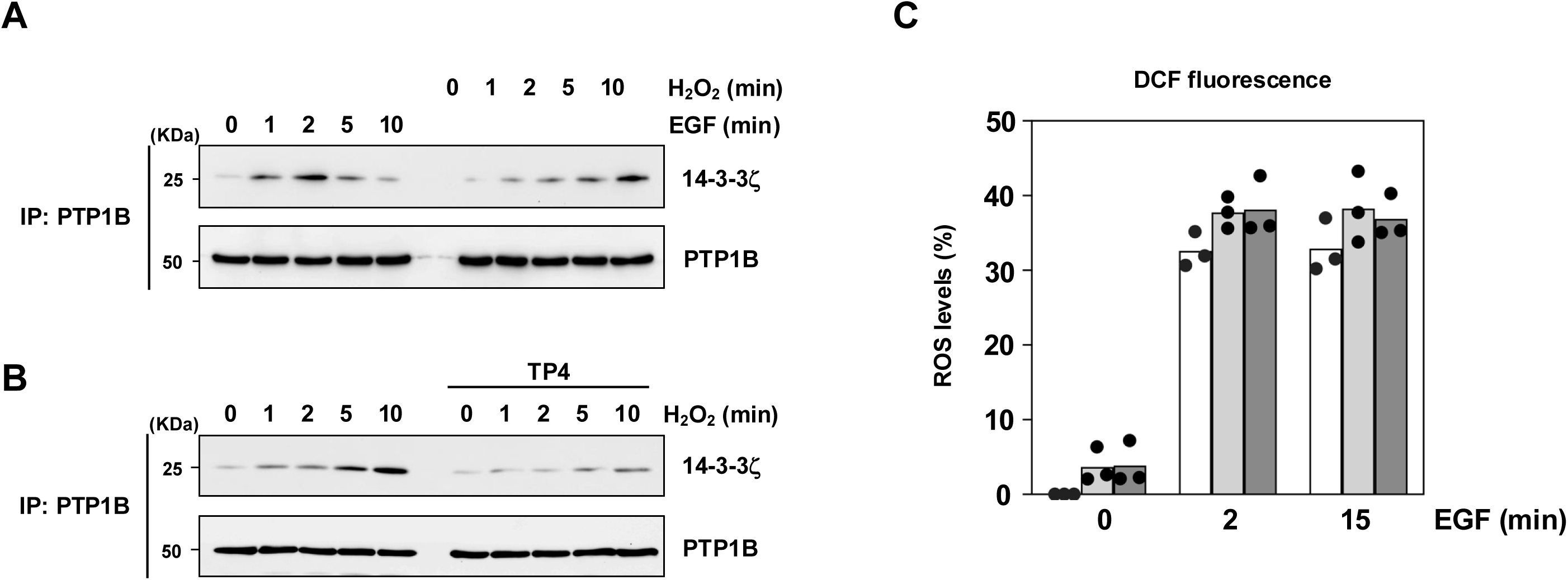
TP4 disrupts H_2_O_2_-mediated association of PTP1B and 14-3-3ζ. (*A*) HEK293T cells were stimulated with EGF or H_2_O_2_ for the indicated times. PTP1B was immunoprecipitated and proteins were separated by SDS-PAGE and blotted for 14-3-3ζ. Lysates were blotted for 14-3-3ζ and PTP1B to control for protein expression. This experiment was repeated three independent times with representative data shown. (*B*) PTP1B was immunoprecipitated from HEK293T cells exposed or not to TP4, stimulated with H_2_O_2_ for the indicated times. Immunoprecipitated proteins and lysates were separated by SDS-PAGE and blotted for 14-3-3ζ and PTP1B. This experiment was repeated three independent times with representative data shown. (*C*) Levels of EGF-mediated H_2_O_2_ production were quantitated by DCF fluorescence in HEK293 cells pretreated or not with TP4. The average readout value of technical replicates is represented by the dot-plot bar graph. N = 3 independent experiments are shown.

### TP4-Mediated Activation of PTP1B Leads to EGFR Dephosphorylation and Decreased Effectiveness of Cancer Cells to Form Colonies

Since EGFR is a PTP1B substrate, we examined whether keeping PTP1B active with TP4 affected EGFR phosphorylation. Immunoblots for phospho-Tyr^992^ and Tyr^1148^, two known PTP1B sites (24), showed that TP4 pretreatment before EGF stimulation markedly reduced phosphorylation at these sites (Fig. 4A). In contrast, phosphorylation of Tyr^845^, a SRC-specific site (25), was strongly increased in TP4-pretreated EGF-stimulated 293 cells, consistent with enhanced PTP1B-mediated SRC activation (7). Alternatively, measuring total EGFR tyrosine phosphorylation levels did not show any overall changes (Fig. S4A). We then assessed whether PTP1B reactivation could prevent colony formation of A431 cells, EGFR-driven epidermal cancer cells in which PTP1B is reversibly oxidized (9, 26). Exposing A431 epidermoid carcinoma cells to TP4 led to marked inhibition of colony formation and cell viability consistent with a tumor suppressive role of PTP1B in this context (Fig. 4B, C and Fig. S3B). Importantly, TP4 also showed similar effects on the dephosphorylation of Tyr^992^ and Tyr^1148^ in A431 cells (Fig. S3C). These data highlight the ability of phosphatases to downregulate important signaling pathways in cancer cells when activated using small molecules such as our protein-protein interaction inhibitor that activates PTP1B.

**Fig. 4.**
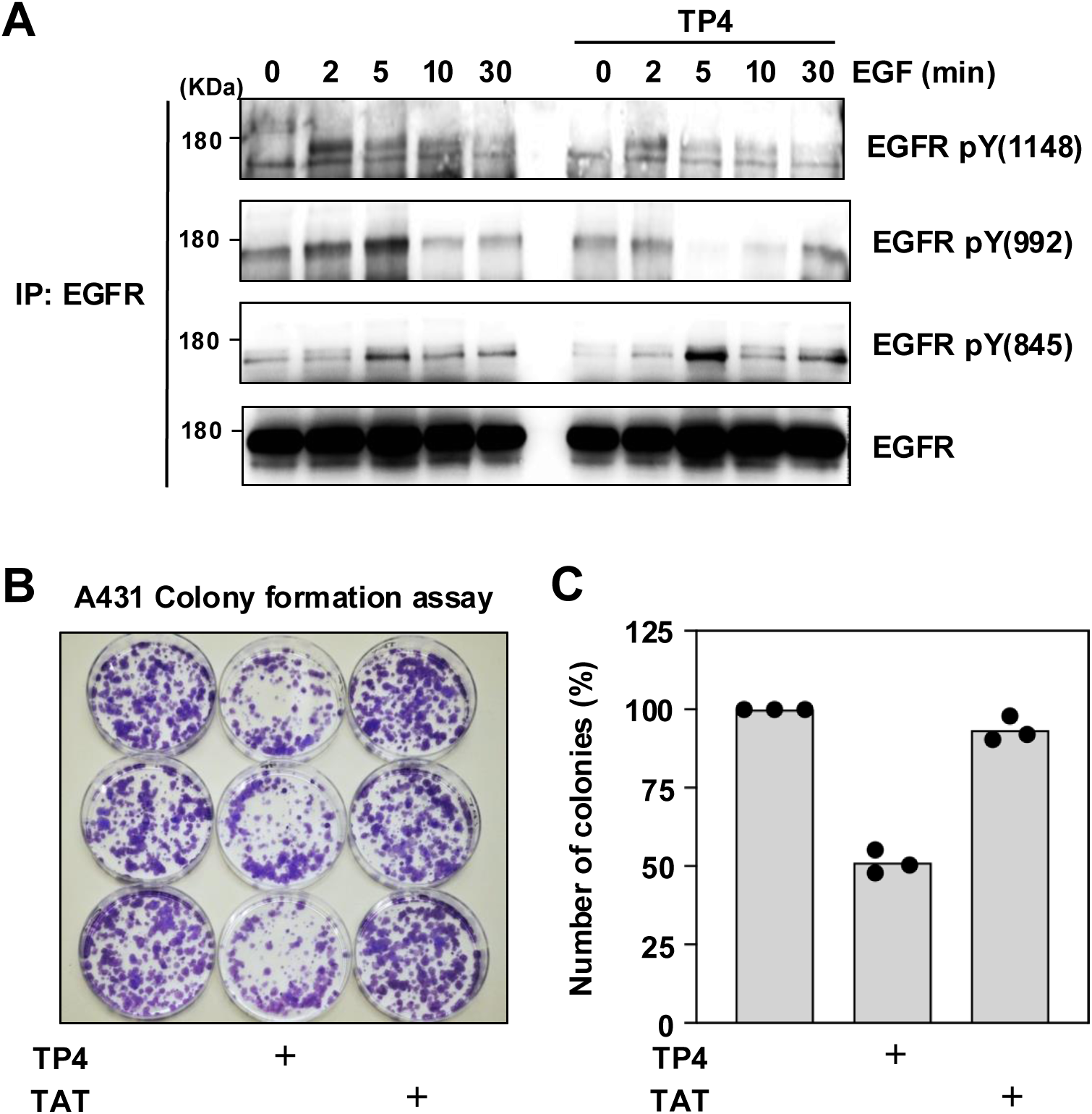
Activating PTP1B using TP4 decreases phosphorylation of specific EGFR sites and prevents colony formation. (*A*) EGFR was immunoprecipitated from HEK293T cells treated or not with TP4 and stimulated with EGF for the indicated times. Proteins were resolved on SDS-PAGE and EGFR phosphorylation was assessed by immunoblot using anti-phospho pY^1148^, anti-phospho pY^992^, anti-phospho pY^845^ antibodies. This experiment was repeated three independent times with representative data shown. (*B*) Inhibition of A431 colony formation by TP4 was assessed by exposing A431 cells to TP4 or TAT peptides for 12 days. Cells were fixed and stained with crystal violet for visualization. (*C*) Quantitative analysis of A431 colony formation in presence of TAT alone, or TP4. The average readout value of technical replicates for control-, TP4- and TAT-treated cells at day 12 is represented by the dot-plot bar graph. N = 3 independent experiments are shown.

### Unphosphorylated and Phosphorylated Forms of the PTP1B Activation Peptide Decrease EGFR Phosphorylation

Results from our TAT-Peptide 4 studies showed that the TAT moiety grants cell permeability to the pTyr loop-derived peptides to activate PTP1B. We next wanted to enquire whether the phosphorylation state of our pTyr loop-derived peptide was important to activate PTP1B. However, TAT has shown limited effectiveness in facilitating cell permeability of phosphorylated peptides. To circumvent this issue, we used pHLIP [pH(low)insertion peptide], a molecule that recognizes the acidic environment of cancer cells to insert itself into their plasma membrane and deliver a cargo peptide to their intracellular milieu via a cleavable disulfide bond (Fig. 5A) (27, 28). We first generated an unphosphorylated version [pHLIP-Peptide 4 (pHLIP-P4)], and a Ser^50^-phosphorylated version (pHLIP-phosphoP4) of our PTP1B activating peptide bound to pHLIP and confirmed their ability to insert themselves into liposomes at low pH by circular dichroism spectroscopy (Fig. S4A,B) and by tryptophan fluorescence (Fig. S4C,D). We then pre-treated A431 cells with pHLIP-P4 at pH 6, followed by EGF exposure for the indicated times. In these conditions pHLIP-P4 could indeed decrease EGFR phosphorylation on the Tyr^992^ and Tyr^1148^ sites (Fig. 5B, Fig. S5A), as seen with the TP4 peptide in Figures 4A and S3C. Interestingly, decreased EGFR phosphorylation was also observed when cells were preincubated with the phosphorylated pTyr loop-derived peptide on Ser^50^ (pHLIP-phosphoP4) (Fig. 5C, Fig. S5B). As expected from previous studies showing that pHLIP did not get inserted into membranes and deliver its cargo into cells at pH 8 (27, 28), pre-treating A431 cells with pHLIP-P4 or pHLIP-phosphoP4 at pH 8 did not affect EGFR phosphorylation (Fig. 5D, E and Fig. S5C, D). Overall, our findings indicate that disrupting the PTP1B-14-3-3ζ-complex with tailored therapeutic molecules represents a promising strategy to constrain hyperactive growth-factor signalling pathways involving PTP1B (Fig. 6). In addition, using pHLIP to deliver PPI inhibitors that activate PTP1B may provide a strategy to counteract dysregulated signaling networks within the acidic tumor microenvironment, while preventing unwanted side effects of activating PTP1B in healthy tissues.

**Fig. 5.**
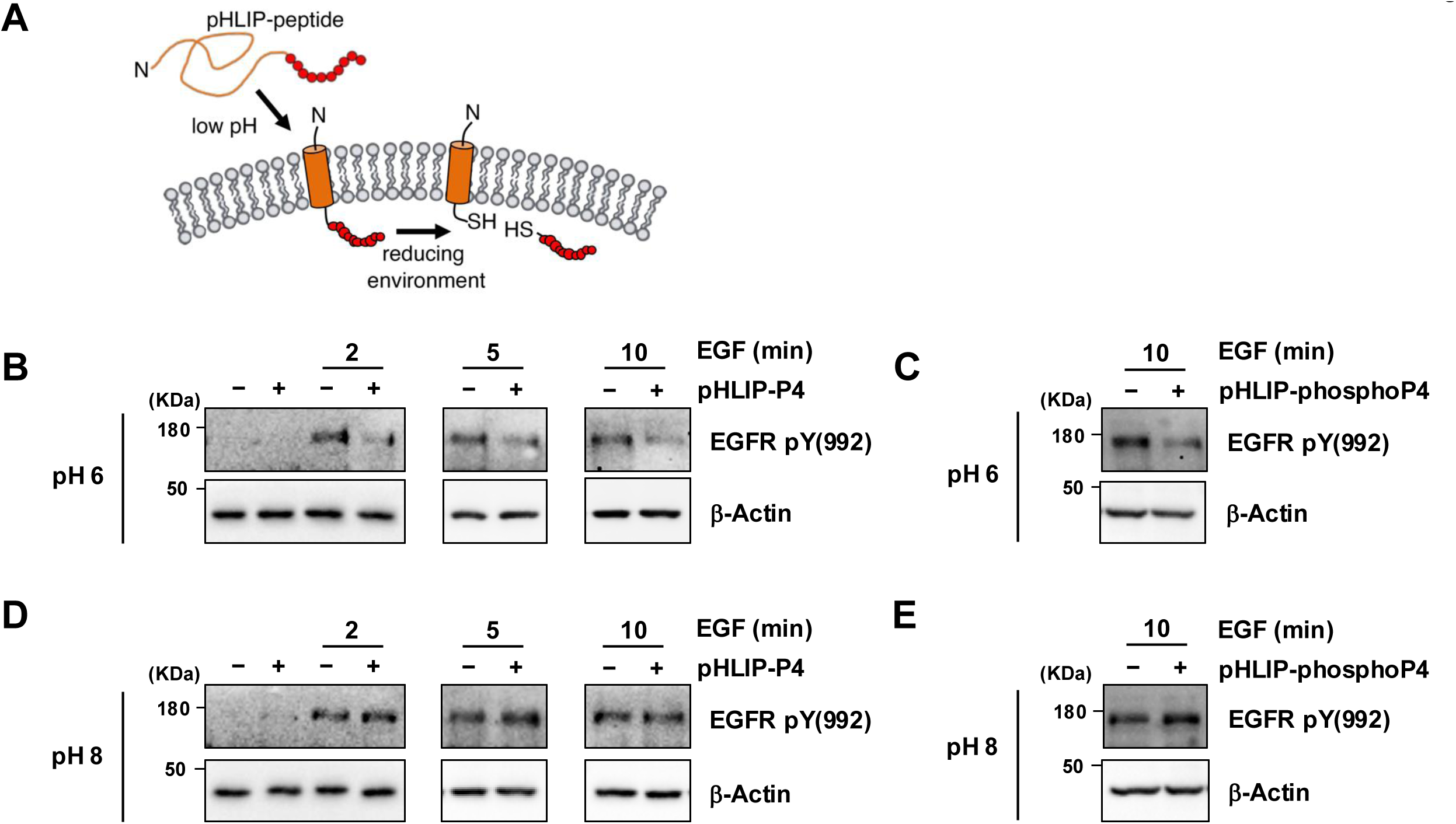
pHLIP-P4 reduces EGFR phosphorylation in a pH-dependent manner. (*A*) Schematic of the mechanism of action of the pH(low) insertion peptide (pHLIP), a molecule that can selectively introduce peptides into cancer cells based on their extracellular acidic environment. A pHLIP molecule (orange sequence) is covalently joined to a 10 amino acid-long PTP1B activation peptide (P4, red circles) via a disulfide bond. Lowering the culture media to pH 6 allows pHLIP to change conformation and insert itself into the membrane of A431 cells. Following this insertion, the reducing intracellular environment leads to the reduction of the pHLIP-P4 disulfide bond and the release of P4 in the intracellular milieu, near EGFR receptors. A431 cells, cultured at pH 6, were treated with pHLIP-P4 (*B*) or pHLIP-phosphoP4 (*C*) followed by EGF treatment for the indicated times. Proteins were resolved on SDS-PAGE and EGFR phosphorylation was assessed by immunoblot using anti-phospho pY^992^ and protein loading was estimated by probing for β-actin. A431 cells, cultured at pH 8, were treated with pHLIP-P4 (*D*) or pHLIP-phosphoP4 (*E*) followed by EGF treatment for the indicated times. Proteins were resolved on SDS-PAGE and EGFR phosphorylation was assessed by immunoblot using anti-phospho pY^992^ and protein loading was estimated by probing for β-actin. These experiments were repeated three independent times with representative data shown.

**Fig. 6.**
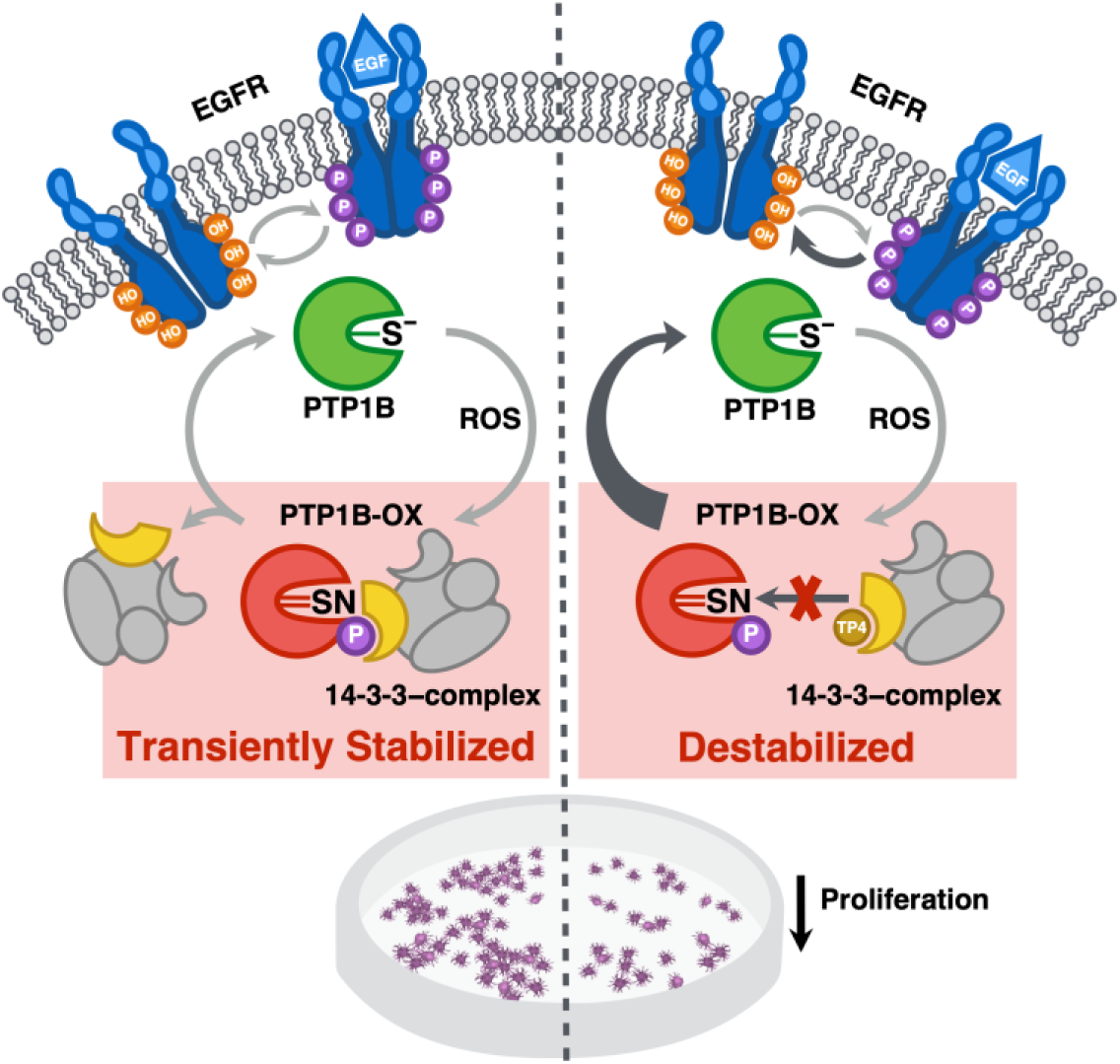
Activation of PTP1B using a protein-protein interaction inhibitor in cells. In resting cells, PTP1B is active and possesses a reactive catalytic cysteine residue (S^−^). Following EGFR activation, a ROS-producing stimulus, PTP1B becomes rapidly and transiently inactivated by ROS. Key conformational changes of the inactive sulphenyl-amide (SN) species of PTP1B-OX include one element of the active site, the phospho-tyrosine recognition loop containing the sequence Lys^41^ to Ser^50^. Upon PTP1B oxidation, this loop adopts a cytosol exposed position which leads to Ser^50^ phosphorylation and binding to a 14-3-3ζ–complex. In turn, binding of 14-3-3ζ–complex to PTP1B-OX transiently stabilizes the inactive, oxidized form of the enzyme and allows phosphorylation of its substrates. On the other hand, preventing the transient interaction between 14-3-3ζ–complex and PTP1B-OX using TP4, perturbed the redox cycle of PTP1B and effectively activated PTP1B. PTP1B activation by TP4 decreased EGFR phosphorylation and inhibited the proliferation of EGFR-driven human cancer cells.

## Discussion

Four decades of research has shed light on the complex regulation of PTPs by post translational modifications (7). Several of these studies established that although PTPs could be oxidized by a plethora of ROS, very few ROS appeared to be involved in redox signaling *in vivo* (3, 6, 29). Our group and others have previously demonstrated that dismutation of superoxide into H_2_O_2_, and the subsequent reaction of H_2_O_2_ with bicarbonate to form peroxymonocarbonate (HCO_4_^−^) were essential steps leading to the oxidation of PTPs in cells (4, 30–32). In this context, the prevailing view is that cellular oxidants generated nearby RTKs diffuse through surrounding antioxidant barriers and eventually reach PTPs to selectively oxidize their low-p*K*a catalytic cysteine residue. Hence, it is thought that a redox network regulating PTP inactivation and reactivation exerts control and fine-tunes protein phosphorylation (17). Since sustained phosphorylation is a critical determinant of normal *versus* pathological signaling, developing tyrosine kinase inhibitors or pharmacological activators of PTP is key to restoring physiological signaling. At this point, although several PTP inhibitors show promise (7, 33), no compounds capable of specifically activating PTPs has been reported. We describe here for the first time the design of a protein-protein interaction inhibitor that enabled us to activate PTP1B. Our data highlights the importance of destabilizing the assembly of a PTP1B-multiprotein complex that regulates the oxidation of PTP1B to maintain PTP1B in its active form and decrease phosphorylation of its substrates as a potential therapeutic strategy.

Two loops of PTP1B become exposed to the cytosol when the enzyme is reversibly oxidized [Fig. S1,(15, 16)]. Interestingly, these structural changes can be taken advantage of by conformation sensor antibodies to lock PTP1B-OX into its inactive form (34). Similarly, we have previously shown that 14-3-3ζ stabilizes the reversibly oxidized form of PTP1B by associating with one of these two loops, the pTyr loop, in cells (18). Since 14-3-3s are phosphoserine binding proteins, it is not surprising that our *in vitro* pulldown experiment showed that Ser^50^ phosphorylation was essential for 14-3-3ζ to associate with the pTyr loop of PTP1B. This also confirmed our previous surface plasmon resonance results that showed phosphorylation of PTP1B-OX to be essential for 14-3-3ζ binding (18). However, pulldowns from lysates are more nuanced, as the non-phosphorylated version of the pTyr-derived peptide also demonstrated measurable binding to 14-3-3ζ. These findings indicate that while 14-3-3ζ interacts with phosphorylated PTP1B-OX, this interaction likely occurs within a larger multiprotein complex that also associates with the unphosphorylated form of the enzyme. This interpretation is further supported by our results showing that TAT-peptides lacking Ser^50^ in their sequence also inhibit the association of 14-3-3ζ with PTP1B following EGFR activation. Given that the pTyr loop resides on the surface of PTP1B at the periphery of the catalytic pocket when the enzyme is reduced, it is plausible that a component of the 14-3-3ζ-associated multiprotein complex engages this loop in the reduced state. Since inhibiting this interaction keeps PTP1B in its reduced form, we hypothesize that a component of this complex engages with the pTyr loop to facilitate oxidation transfer onto the adjacent catalytic cysteine residue.

The ability of PTP1B to regulate signaling is highly dependent on its subcellular localization. PTP1B resides on the cytoplasmic face of the endoplasmic reticulum (ER), anchored by a small C-terminal sequence (35), and it is from its ER-bound location that PTP1B dephosphorylates EGFR following ligand stimulation (36). Dephosphorylation of the endocytosed EGFR occurs at direct membrane contact sites between endosomes and the ER, an area that provides sites of protein-protein interaction between organelles in what has been referred to as a potential “dephosphorylation compartment” (34, 37). These restricted contact points between PTP1B and EGFR enable signaling by both plasmalemmal and cytosolic EGFR and help regulate the duration of EGF signaling. (38). This also appears to be the case for other RTKs such as the insulin receptor or Met (39, 40). Thus, the pronounced dephosphorylation of PTP1B-regulated sites on EGFR at 5 and 10 minutes after EGF stimulation in TP4-treated cells likely reflects the association of endosomes with the ER and subsequent dephosphorylation by ER-anchored PTP1B. Moreover, in response to EGF, PTP1B oxidation is also regulated in a spatially dependent manner by NOX4, another ER-resident protein (41). Interestingly, although H_2_O_2_ can readily diffuse, a cytosolic form of PTP1B that do not encode its ER-anchoring sequence is not regulated by NOX4 (41). Hence, similar to its localized regulation of EGFR dephosphorylation, PTP1B’s catalytic activity is also controlled in a location-dependent fashion. Taken together, our findings show that the PPI preserves PTP1B activity despite its spatial proximity to NOX4.

NOX4 has been reported to exert both pro- and antiproliferative effects depending on the cellular context, highlighting its complex role in ROS signaling (41–44). Although early studies questioned its proliferative function in NIH3T3 cells, several loss-of-function studies demonstrate that NOX4 depletion reduces proliferation in multiple cancer cell types (45, 46). When considering PTP1B as a downstream effector of NOX4, knocking down NOX4, or decreasing its activity, would likely lead to increased PTP1B activity and decreased phosphorylation of its substrates. Hence it is not surprising that our results in A431 cancer cells exposed to TP4 show decreased effectiveness of cancer cells to form colonies, consistent with these reports.

With more than eighty approved peptide drugs, their higher clinical trial success rates and ability to target ‘undruggable’ PPIs, peptides have carved out an important niche in drug development, complementing both small-molecule and biologic therapeutics (47, 48). Although small-molecule drugs offer advantages such as low cost, oral bioavailability, and efficient membrane permeability, their small size limits their ability to effectively inhibit large protein-protein interaction interfaces (49, 50). Because of their size, small molecules usually contact only 300 to 1000 Å² of a protein surface, whereas protein-protein interactions typically involve considerably larger interfaces ranging from 1500 to 3000 Å² (50). Accordingly, in contrast to small molecules, the larger size, conformational flexibility and enhanced specificity of peptides allow them to engage and inhibit the broad interfaces typical of PPIs (47). Following the identification of promising peptide drug candidates, a critical challenge is to address the well-known intrinsic limitations of peptides, particularly their typically low membrane permeability and short *in vivo* half-life (51). Our proof-of-concept experiments showed that TP4, the TAT-PPI version of our peptide, effectively disrupted the interaction between PTP1B and the 14-3-3ζ-complex in cells, thereby activating PTP1B. However, because TAT is a broadly cell-permeable peptide, its use to therapeutically deliver intracellular peptides would likely lead to off-target toxicity, given the ubiquitous expression of PTP1B. As an initial strategy to mitigate potential bioavailability limitations, we functionalized our PPI peptide with the pH(low) insertion peptide (pHLIP). This peptide uniquely enables selective translocation of conjugated cargo into cancer cells and tumors in mice by exploiting their acidic extracellular environment, while minimizing uptake in healthy tissues (52–54). Importantly, unlike TAT, pHLIP transports its cargo by forming a transmembrane α-helix that inserts across the lipid bilayer in acidic environments. In this process, the C-terminus enters the cytoplasm with its cargo while the N-terminus remains extracellular (55). Since most solid tumors exhibit elevated extracellular acidosis regardless of their origin, the acidic microenvironment of cancer cells (pH 6.0–6.8) provides a broadly applicable targeting mechanism that both increases bioavailability and decreases resistance (56). Using pHLIP as a delivery method, we showed that our PTP1B activating peptide can get delivered in cancer cells at pH 6 and decrease the phosphorylation of PTP1B sites on EGFR. The signaling changes observed in these experiments suggest that the delivery of our PPI peptide activates PTP1B in these cells, leading to decreased EGFR signaling in epidermoid cancer cells.

Until now, no selective peptide or small-molecule activator of PTP1B has been identified. Thus, developing a protein-protein interaction inhibitor that activates PTP1B by blocking its association with the 14-3-3ζ complex constitutes a significant conceptual and mechanistic advance. Overall, our data revealed that activating PTP1B using a PPI inhibitor was an effective approach to fine-tune EGFR signaling and limit the survival of cancer cells. Our findings provide new insight into the redox regulation of PTP1B and demonstrate that modulating the redox state of PTPs with PPI inhibitors constitutes a promising strategy for developing small-molecule or peptide-based agents capable of activating specific PTPs *in vivo* and restoring physiological signaling.

## Materials and Methods

### Material

Anti-EGFR and 14-3-3ζ antibodies were from Santa Cruz Biotechnology. Anti phosphoEGFR antibodies were from Cell Signaling Technology. HA-peroxidase was from Roche. PT-66-aga e-conjugated beads, anti-FLAG M2 beads, and anti-HA beads and anti-Flag M2 peroxidase were purchased from Sigma. His-tagged 14-3-3ζ, β-actin were from Abcam. Streptavidin-Sepharose and streptavidin-HRP was from Cytiva. HRP-conjugated secondary antibodies were from Jackson ImmunoResearch Laboratories, Inc. Protease inhibitor mixture tablets were from Roche. Catalase and superoxide dismutase were from Calbiochem. Surfact-Amps Nonidet P-40, zeba desalt spin columns, EZ-Link biotin-iodoacetyl-PEG2 (biotin-IAP), and iodoacetic acid were from ThermoScientific. pTyr loop-derived peptide (CKNRNRYRDVS), phosphoSer^50^ pTyr loop-derived peptide (C<u>KNRNRYRDVpS),</u> TAT (GRKKRRQRRRPQ) and TAT-<u>pTyr loop-derived peptides</u> (TP1-4: TP1: <u>KNRNRYRDVS-</u>GRKKRRQRRRPQ; TP2: <u>KNRNRYRDV-</u>GRKKRRQRRRPQ; TP3: GRKKRRQRRRPQ-<u>KNRNRYRDV</u>; TP4: GRKKRRQRRRPQ-<u>KNRNRYRDVS</u>) were from GenScript USA Inc.

### Cell culture

HEK293T cells were maintained in culture at 40-90% confluency in EMEM (Eagle’s Minimum Essential Medium, 1000 mg/l glucose, ATCC) containing 100 U/ml penicillin, 100 μg/ml streptomycin and 10% FBS at 5% CO_2_ and 37°C. 4 μg of plasmid DNA was routinely transfected using TurboFect (ThermoFisher) on 80% confluent cells. Transient expression was allowed to progress for 48 hours. Cells were serum starved overnight using EMEM without serum. A431 cells were maintained in DMEM supplemented with 100 U/ml penicillin, 100 μg/ml streptomycin and 10% FBS at 5% CO_2_ and 37°C.

### pTyr loop derived-peptide pulldowns

HEK293 cells were lysed in a lysis buffer consisting of 20 mM HEPES pH 7,4, 150 mM NaCl, 1% NP40, 1 mM EDTA, 10 mM NaF, 25 µg/ml aprotinin, 25 µg/ml leupeptin, 1 µM microcystin and 100 nM okadaic acid. The protein content of lysates was measured by Bradford assay, and equal amounts of lysates were incubated with the pTyr loop-derived peptide (C-KNRNRYRDVS), a phospho-Ser^50^ pTyr loop-derived peptide (C-KNRNRYRDVpS) or L-Cys coupled to BcMag Iodoacetyl-activated magnetic beads (BioClone Inc.) for 3 hours at 4 °C. The beads were washed on a magnetic stand three times with lysis buffer. The beads were then resuspended in 20 μl of 4X Laemmli sample buffer and heated at 95°C for 2 minutes. Proteins were separated by SDS-PAGE and detected by immunoblotting. The interaction between the phospho-Ser^50^ pTyr loop-derived peptides and 14-3-3ζ was performed similarly, using purified his-tagged 14-3-3ζ.

### Immunoprecipitation and immunoblotting

FLAG-PTP1B, HA-14-3-3ζ and EGFR were immunoprecipitated as follows. Cells were grown to 80% confluence in 10-cm plates, transfected for 48 hours and serum-starved for 16 hours. Following serum-starvation, cells were stimulated with EGF (100 ng/ml) to activate EGFR for the indicated times. After treatment, the plates were transferred on ice, washed with cold PBS and extracted in 800 μl of a lysis buffer consisting of 20 mM HEPES pH 7,4, 150 mM NaCl, 1% NP40, 1 mM EDTA, 10 mM NaF, 25 μg/ml aprotinin, 25 μg/ml leupeptin, 1 μM microcystin, 100 nM okadaic acid. 1 mM Na_3_VO_4_ was also supplemented to the lysis buffer in experiments assessing EGFR phosphorylation. All subsequent steps were carried out on ice or at 4 °C. Cells were lysed on a rotating wheel at 4°C for 30 minutes. Cell debris were centrifuged at 14,000 x g for 10 minutes, and protein concentrations were determined. 500 μg of protein was diluted in cold lysis buffer and precleared for 20 minutes with sepharose beads. The supernatants were incubated for 3 hours on a rotating wheel with appropriate beads precoupled to -HA, -FLAG, anti-EGFR antibodies or to anti-pTyr antibodies (PT-66). The immune complexes were pelleted at 3000 x *g* for 5 minutes and washed three times with cold lysis buffer. The beads were resuspended in 20 μl of 4X Laemmli sample buffer and heated at 95°C for 2 minutes. Proteins were separated by SDS-PAGE and detected by immunoblotting.

### Assay of PTP oxidation

The cysteinyl-labeling assay was performed as described (5, 21). In brief, cells were starved for 16 hours in serum-free EMEM. For EGF stimulations, cells were stimulated with 100 ng/ml EGF for indicated times and lysed in degassed lysis buffer (50 mM Sodium Acetate (pH 5.5), 150 mM NaCl, 1% NP40, 10% (v/v) glycerol) supplemented with 10 μg/ml aprotinin, 10 μg/ml leupeptin, 10 mM IAA, 250 U/ml catalase and 125 U/ml superoxide dismutase after which alkylation was allowed for 1hour at room temperature. Lysates were cleared by centrifugation at 10,000 rpm for 10 minutes, and buffer exchanged with (1 mM) TCEP-containing lysis buffer using Zeba columns (Pierce). Lysates were reduced for 30 minutes and supplemented with (5 mM) a biotin-labeled iodoacetic acid probe (Pierce). Labeled PTP1B was pulled down with streptavidin-Sepharose beads, boiled for 2 minutes in sample buffer and used for immunoblotting.

### Assessment of ROS production in cells

ROS production was estimated by using 2′,7′-dichlorofluorescein diacetate (DCF-DA) dye by measuring the conversion of non-fluorescent DCF-DA to fluorescent dichlorofluorescein (DCF) within the cell. Briefly, cells were seeded in black 96-well tissue culture plates at a density of 10,000 cells/well. Next day the cells were deprived of serum for 16 hours following which they were washed with pre-warmed HBSS (calcium and magnesium free). Cells were incubated with 10 μM DCF-DA for 30 minutes in serum- and phenol red-free media at 37°C and washed again with pre-warmed HBSS. This was followed by incubation with indicated treatments of TAT or TP4 for 1 hour and EGF for the indicated times. The measurement of intracellular ROS was carried out at 485 nm excitation and 535 nm emission wavelengths after each tested time.

### Immune-complex PTP activity assay

Measurement of endogenous PTP1B activity was assessed in HEK293T cells as previously described (20). 60-80% confluent cells were transfected with PTP1B-FLAG and 14-3-3ζ**-**HA for 48 hours. Cells were serum-starved 32 hours post-transfection for a 16-hour period and stimulated with EGF (100 ng/ml) for the indicated times, in presence, or absence of TP4 (15 μM, 1 hour). Cells were lysed under strict hypoxic conditions, on ice following with a cold degassed lysis buffer consisting of 25 mM HEPES pH 7.4, 150 mM NaCl, 1 mM EDTA and 1% NP-40 (Thermo Fisher). Degassing of the buffer is a critical step for this assay. Cells were lysed on a rotating wheel at 4°C for 30 minutes. Cell debris were centrifuged at 14,000 x *g* for 10 minutes, and protein concentrations were determined. The supernatants (200 μg) were incubated at 4°C for 3 hours on a rotating wheel with 10 μl anti-FLAG Dynabeads (Life technologies). The immune complexes were then washed with degassed lysis buffer and resuspended in pNPP assay buffer (20 mM HEPES pH 7.4, 100 mM NaCl and 0.05% w/v BSA, 20 mM pNPP), with or without DTT (5 μM). The beads were protected from light, placed on a rotator at room temperature, and the converted substrate was measured following a 30 min incubation. 80 μl of the supernatant from the enzymatic reaction was stopped with 20 μl of 2M NaOH and absorbance was measured at 405 nm using a spectrophotometer (SpectraMax, Molecular Devices).

### Biotin labeling of active PTP1B

The direct cysteinyl-labeling assay was performed as previously described (20). Cells were starved for 16 hours in serum-free EMEM. For EGF stimulations, cells were stimulated with 100 ng/ml EGF for indicated times and lysed in degassed lysis buffer (50 mM Sodium Acetate (pH 5.5), 150 mM NaCl, 1% NP40, 10% (v/v) glycerol) supplemented with 25 μg/ml aprotinin, 25 μg/ml leupeptin and 10 mM IAP under strict hypoxic conditions. Alkylation of active PTPs with biotin-IAP was performed for 1 hour at room temperature. Lysates were cleared by centrifugation at 10,000 rpm for 10 minutes, and biotin IAP-labeled PTP1B was pulled down with streptavidin-Sepharose beads, boiled for 2 minutes in sample buffer and used for immunoblotting.

### Colony formation assay

A431 cells were seeded (1500 cells) in 60 mm petri dishes in 3 ml DMEM supplemented with 10% FBS. After 24 hours of seeding, cells were either treated with 15 μM of TP4 or TAT peptide or left untreated. Cell media was replenished, with their respective treatment, every day for a 12-day period. After 12 days, media was carefully removed and colonies were fixed with 100% methanol for 20 minutes, washed with ddH_2_O and stained with a 0.5% crystal violet solution for 5 minutes. The colonies were rinsed to remove excess dye, dried overnight and counted manually under microscope. Clones of 50 or more cells counted as one colony.

### Cell viability assay

3-(4,5-demethylthiazol-2-yl)-2,5-diphenyltetrazolium bromide (MTT) conversion to formazan is an indicator of mitochondrial dehydrogenase activity of live cells, which is inferred as a measure of percent cell viability. A431 cells were seeded as described in colony formation assay. After 24 hours of plating, cells were either treated with 15 µM of TP4, TAT peptide or left untreated. Media was replaced along with their respective treatments every day for 5 days. MTT (Sigma Aldrich) was added (300 μl/plate of 5 mg/ml stock solution) 3 hours prior to completion of the test time. After carefully aspiration of media from the plates, 1.5 ml of DMSO/plate was added for solubilization of formazan crystals. After 10 minutes of incubation, absorbance was measured at 570 nm.

### Peptide synthesis and conjugation

pHLIP, containing a cysteine residue at its C-terminus (GGEQNPIYWARYADWLFTTPLLLLDLALL VDADEGT<u>C</u>G), was prepared by Fmoc solid-phase peptide synthesis using Rink Amide resin on a CEM Liberty BlueTM microwave peptide synthesizer. pHLIP and peptide conjugates were purified by reverse-phase high-performance liquid chromatography (Phenomenex Luna Omega, 5 μm, 250 x 21.20 mm C18; flow rate 5 ml/min; phase A: water 0.1 % TFA; phase B: acetonitrile 0.1 % TFA; 60 min gradient from 95 : 5 to 0 : 100 A/B), and identity was confirmed by matrix-assisted laser desorption ionization time-of-flight (MALDI-TOF) mass spectroscopy (Shimadzu 8020), as previously described (27, 28, 57–59). To prepare the peptide conjugates, phosphoS^50^-P4 and non-pS^50^-P4 peptides were solubilized in water and added in a 1:1 molar ratio to pHLIP (solubilized in DMF/PBS, pH 7.4). The mixture was adjusted to pH 8.4 and incubated for 3 hours at room temperature. The resulting conjugates were again purified by RP-HPLC as described above. The purity of each peptide and conjugate (> 90 %) was determined by analytical RP-HPLC (Phenomenex Nucleosil, 5 μm, 250 × 3.2 mm C18; flow rate 0.5 ml/min; phase A: water 0.01 % TFA; phase B: acetonitrile 0.01 % TFA; 60 min gradient from 95 : 5 to 0 : 100 A/B). Their identity was confirmed by MALDI-TOF MS: pHLIP-P4: purity > 95%; calculated (M + H+) = 5618; found (M + H+) = 5616. pHLIP-phosphoP4: purity > 95%; calculated (M + H+) = 5,698; found (M + H+) = 5,705. The conjugates were quantified at 280 nm by UV/vis absorbance spectroscopy using the molar extinction coefficient of pHLIP (13,940 M^−1^ cm^−1^) and lyophilized.

### Sample preparation of conjugate peptides for circular dichroism (CD) and tryptophan fluorescence measurements

All peptides were first solubilized to 20 μM in 5 mM sodium phosphate (pH 8.0). Each peptide was diluted to a final concentration of 7 μM before analysis. 100 nm 1-palmitoyl-2-oleoyl-sn-glycero-3-phosphocholine (POPC) large unilamelar vesicles were prepared by freeze-thaw and extrusion, as previously reported (27, 28, 57–59). Peptides were incubated with the resulting vesicles at a 1:300 ratio. The pH was adjusted to the desired experimental values with HCl, and the samples were incubated for 30 minutes at room temperature prior to spectroscopic analysis.

### CD spectroscopy

Far-UV CD spectra were recorded on a Jasco J-815 CD spectrometer equipped with a Peltier thermal-controlled cuvette holder (Jasco). Measurements were performed in a 0.1 cm quartz cuvette. Raw data were acquired from 260 to 190 nm at 1 nm intervals with a 100 nm/min scan rate, and at least five scans were averaged for each sample. The spectrum of POPC liposomes was subtracted from all construct samples. CD intensities are expressed in mean residue molar ellipticity [θ] calculated from the following equation:

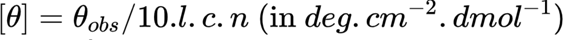

Where *θ_obs_* is the observed ellipticity in millidegrees, *l* is the optical path length in centimeters, *c* is the final molar concentration of the peptides, and *n* is the number of amino acid residues.

### Tryptophan fluorescence spectroscopy

Fluorescence emission spectra were acquired with a Fluorolog-3 Spectrofluorometer (HORIBA). The excitation wavelength was set at 280 nm, and the emission spectrum was measured from 300 to 450 nm with excitation and emission slits set to 5 nm.

### pHLIP-P4 and -phosphoP4 mediated activation of PTP1B

Cells were seeded in 12-well plates at 80,000 cells/well and incubated overnight. The cell media was replaced with serum-free media for 2 hours before peptide treatment. Peptide conjugates were solubilized in an appropriate volume of serum-free media at pH 7.4, so that upon pH adjustment, a concentration of 10 μM was achieved. The peptides were then added to serum-starved cells and incubated for 5 minutes at 37°C. The media was then adjusted to the desired pH using a pre-established volume of serum-free media buffered with citric acid. After a 10-minute incubation at 37°C, the pH media was removed and replaced with fresh serum-free media containing 100 ng/ml EGF, and the cells were incubated for indicated times at 37°C. Following the stimulation period, the media was rapidly removed, and cells were washed in ice-cold PBS. Cells were lysed with 10 mM Tris pH 7.4, 100 mM NaCl, 1 mM EDTA, 1 mM EGTA, 1 mM NaF, 20 mM Na_4_P_2_O_7,_ 2 mM Na_3_VO_4_, 1% Triton X-100, 10% glycerol, 0.1% SDS, 0.5% deoxycholate supplemented with phosphatase and protease inhibitors. Lysates were centrifuged at 10,000 rpm at 4°C for 10 minutes. Sample concentrations were determined and used for immunoblotting.

## Acknowledgments

This research was supported by National Institutes of Health Grants R35GM153449 and R01HL17539101 (to B.B.) and GM139998 (to D.T.). A.B. was a recipient of a fellowship from the Fonds de Recherche du Québec – Santé. BB is also grateful for support from the following foundations: SUNY Research Foundation, Fonds de Recherche du Québec – Santé, Fondation Jacques de Champlain.

## SUPLEMENTARY FIGURES

**Fig. S1.**
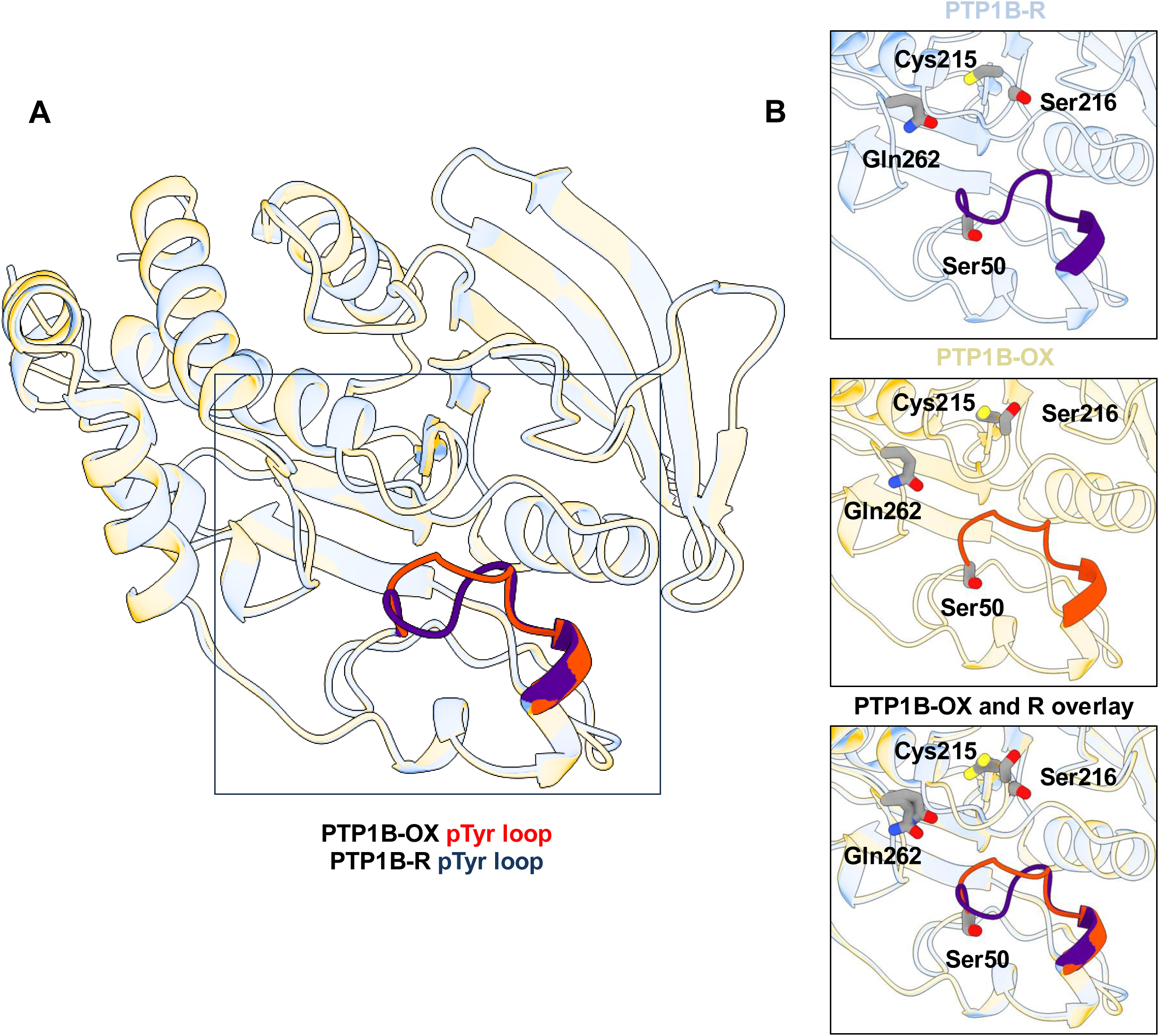
Structure of the reduced and reversibly oxidized form of PTP1B. (*A*) Views of superposed PTP1B-OX (sulfenamide oxidized form, PDB code: 1OEM) and PTP1B-R (reduced form, PDB code:2HNQ). The oxidized PTP1B structure is shown in pale gold while the reduced PTP1B structure is shown in pale blue. The phosphotyrosine binding loop is shown in dark blue for PTP1B-R and in red for PTP1B-OX. (*B*) Close-up views of PTP1B-R, PTP1B1B-OX and superposed view of PTP1B-OX and PTP1B-R.

**Fig. S2.**
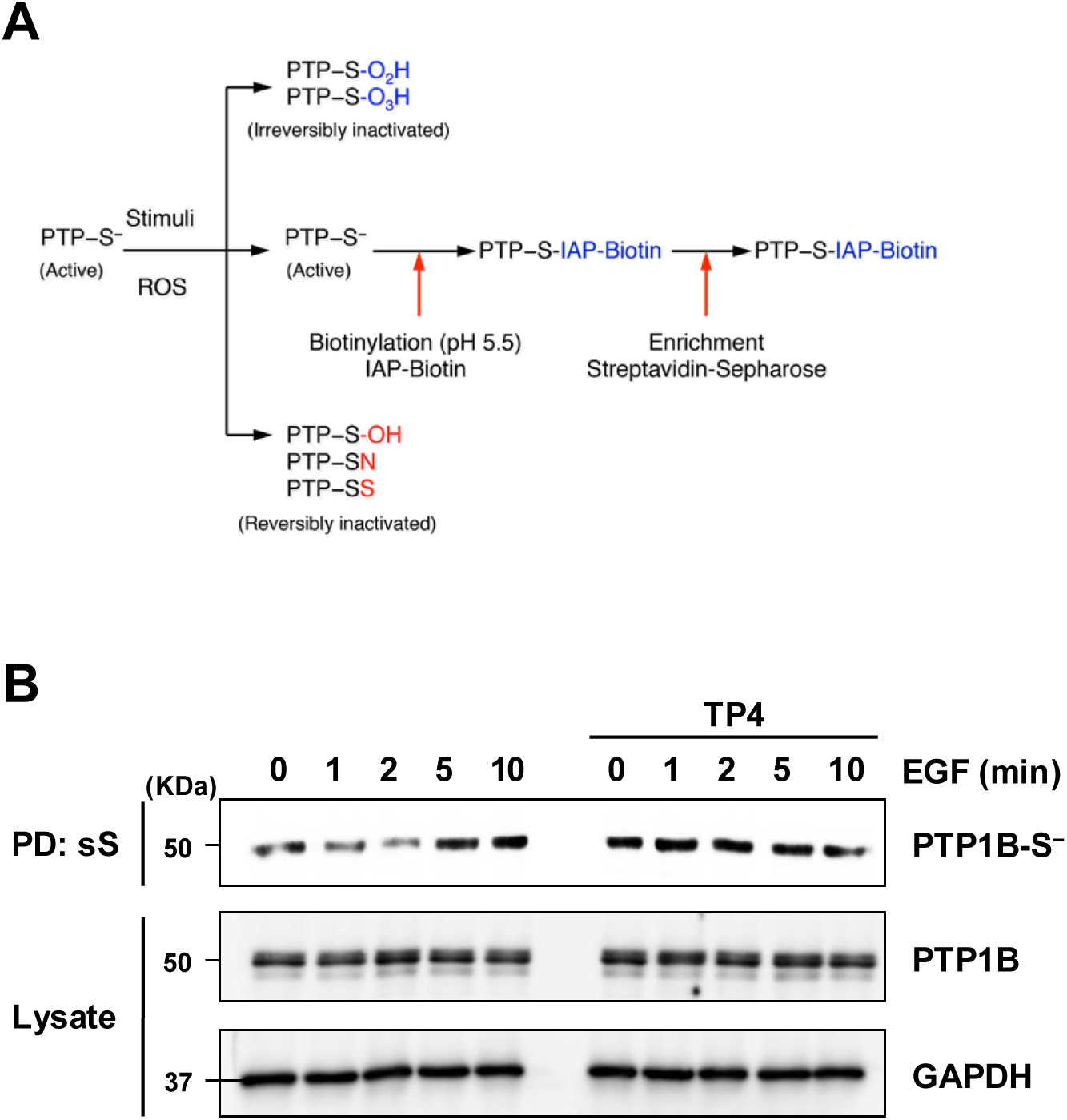
Direct Cysteinyl Labeling of active PTP1B in cell lysates. (*A*) The direct cysteinyl-labeling assay is a 2-step assay designed to assess oxidation of PTPs by labeling active PTPs in cell lysates. In the first step of the assay, the pool of PTP1B that remained active following cell stimulation react and is irreversibly labeled by a biotinylated thiolate-reactive probe (IAP-Biotin) that allows purification by streptavidin pull-down. (*B*) HEK293T cells were pre-incubated or not with TP4, stimulated with EGF for the indicated times, and subjected to a direct cysteinyl-labeling approach. Biotinylated proteins were purified on streptavidin–Sepharose beads, resolved by SDS-PAGE, and blotted for PTP1B. Lysates were probed for PTP1B and GAPDH to control for protein expression and loading. This experiment was repeated three independent times with representative data shown.

**Fig. S3.**
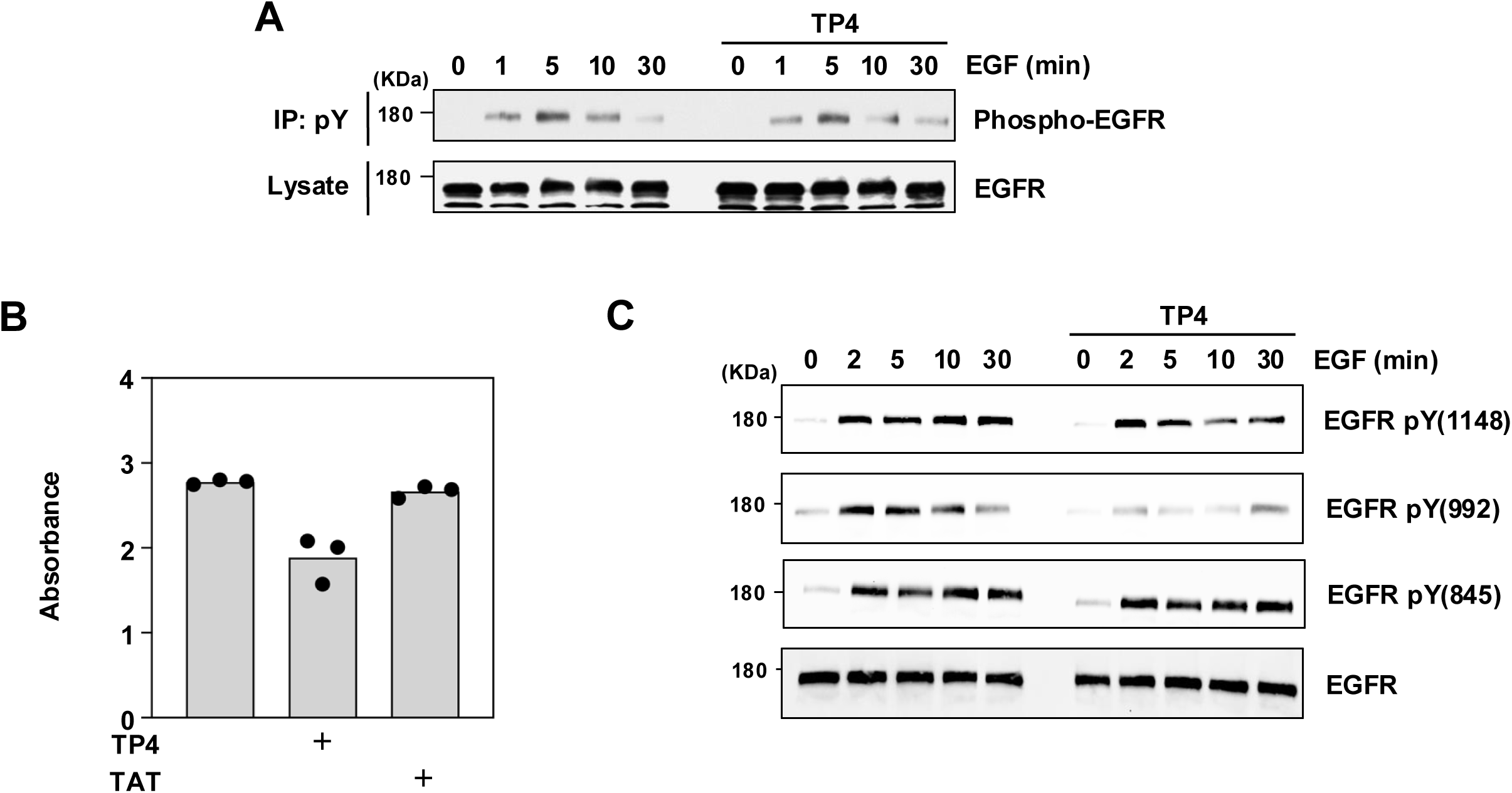
Effects of TP4 on EGFR phosphorylation and cell viability. (*A*) Tyrosine phosphorylated proteins were immunoprecipitated from lysates of serum-deprived HEK293T cells preincubated with TP4 or not (10 μM, 60 min) and stimulated with EGF (100 ng/ml). Proteins were resolved on SDS-PAGE and EGFR phosphorylation was assessed by immunoblot using anti-EGFR antibodies. Lysates were probed for EGFR to control for protein expression. This experiment was repeated three independent times with representative data shown. (*B*) The effect of TP4 on cell viability was assessed by treating A431 cells with regular growth media, or media containing TP4 (15 µM) or TAT (15 µM). The conversion of MTT to formazan was measured at 570 nm following 5 days of treatment. The results show the average absorbance value of technical replicates for control-, TP4- and TAT-treated cells at day 5, represented by the dot-plot bar graph. Absorbance for formazan at 570 nm is a readout for living cells. N = 3 independent experiments are shown. (*C*) A431 cells were treated or not with TP4 and stimulated with EGF for the indicated times. Proteins were resolved on SDS-PAGE and EGFR phosphorylation was assessed by immunoblot using anti-phospho pY^1148^, anti-phospho pY^992^, anti-phospho pY^845^ antibodies. This experiment was repeated three independent times with representative data shown.

**Fig. S4.**
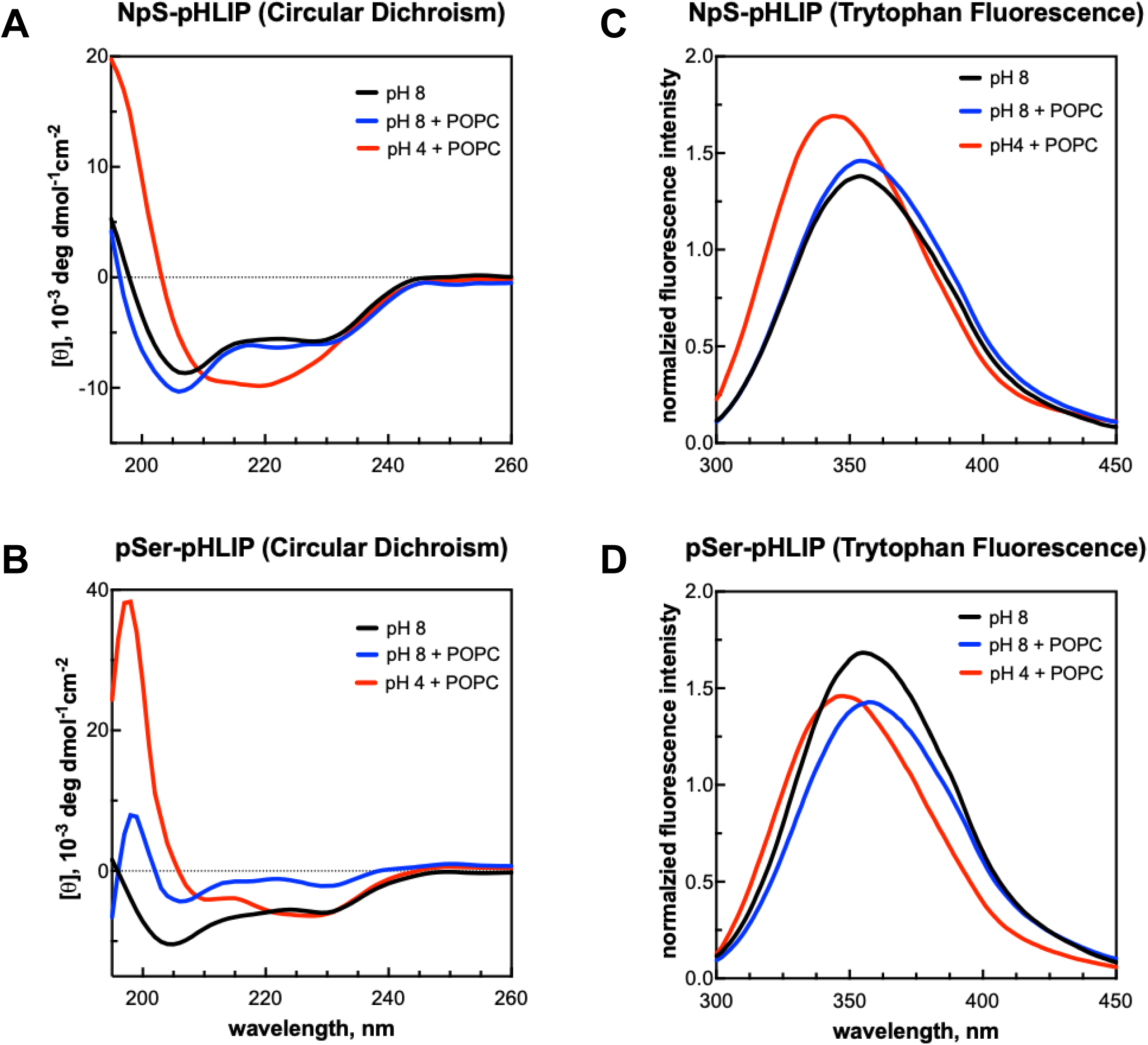
Peptide conjugates insert into lipid vesicle membranes in a pH-dependent manner. The interaction of peptide conjugates with lipid membranes was assessed by circular dichroism (*A*, *B*) and tryptophan fluorescence (*C*, *D*) in the absence (black) or presence of large unilamellar POPC lipid vesicles at pH 8.0 (blue) and pH 6.0 (red).

**Fig. S5.**
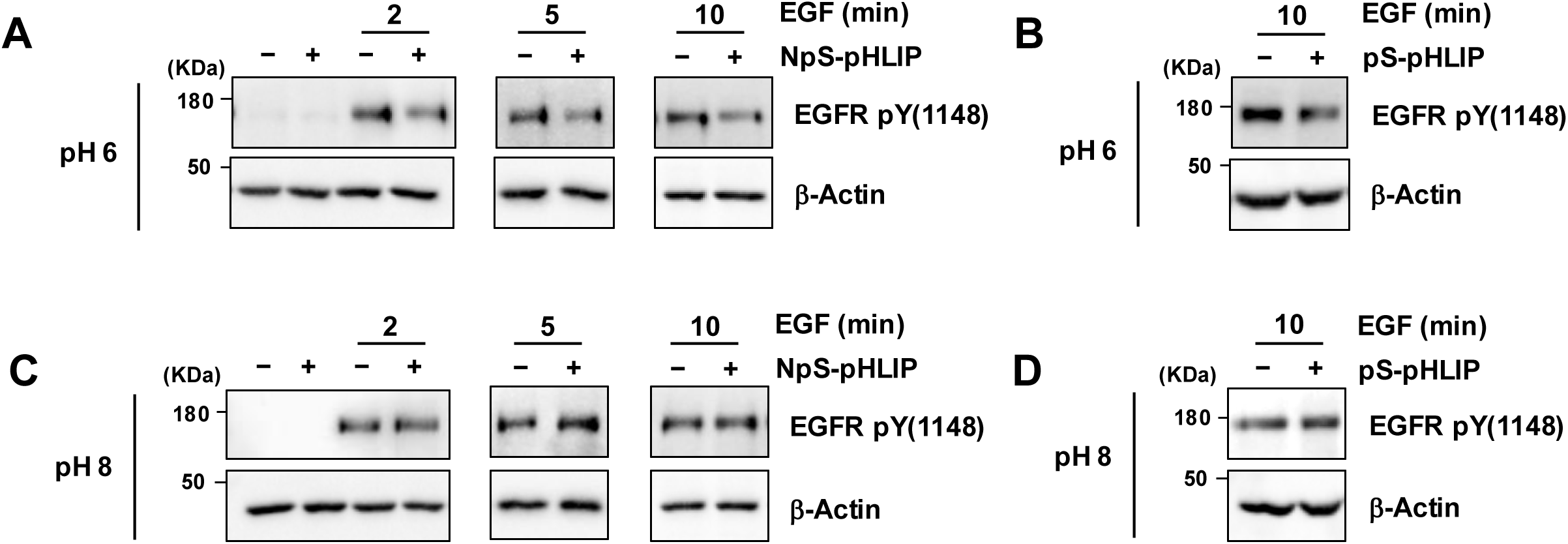
pHLIP-P4 reduces EGFR Tyr^1148^ phosphorylation in a pH-dependent manner. A431 cells, cultured at pH 6, were treated with pHLIP-P4 (*A*) or pHLIP-phosphoP4 (*B*) followed by EGF treatment for the indicated times. Proteins were resolved on SDS-PAGE and EGFR phosphorylation was assessed by immunoblot using anti-phospho pY^1148^ and protein loading was estimated by probing for β-actin. A431 cells, cultured at pH 8, were treated with pHLIP-P4 (*C*) or pHLIP-phosphoP4 (*D*) followed by EGF treatment for the indicated times. Proteins were resolved on SDS-PAGE and EGFR phosphorylation was assessed by immunoblot using anti-phospho pY^1148^ and protein loading was estimated by probing for β-actin. These experiments were repeated three independent times with representative data shown.

## References

1. M. Sundaresan, Z. X. Yu, V. J. Ferrans, K. Irani, T. Finkel, Requirement for generation of H2O2 for platelet-derived growth factor signal transduction. Science 270, 296–299 (1995).

2. H. Sies, D. P. Jones, Reactive oxygen species (ROS) as pleiotropic physiological signalling agents. Nat Rev Mol Cell Biol 21, 363–383 (2020).

3. T. C. Meng, T. Fukada, N. K. Tonks, Reversible oxidation and inactivation of protein tyrosine phosphatases *in vivo*. Mol Cell 9, 387–399 (2002).

4. J. C. Juarez et al., Superoxide dismutase 1 (SOD1) is essential for H2O2-mediated oxidation and inactivation of phosphatases in growth factor signaling. Proc. Natl. Acad. Sci. U.S.A. 105, 7147–7152 (2008).

5. B. Boivin, S. Zhang, J. L. Arbiser, Z. Y. Zhang, N. K. Tonks, A modified cysteinyl-labeling assay reveals reversible oxidation of protein tyrosine phosphatases in angiomyolipoma cells. Proc. Natl. Acad. Sci. U.S.A. 105, 9959–9964 (2008).

6. A. Ostman, J. Frijhoff, A. Sandin, F. D. Böhmer, Regulation of protein tyrosine phosphatases by reversible oxidation. J Biochem. 150, 345–356 (2011).

7. T. Tiganis, N. K. Tonks, Mechanisms, functions and therapeutic targeting of protein tyrosine phosphatases. Nat Rev Mol Cell Biol (2025).

8. K. Loh et al., Reactive oxygen species enhance insulin sensitivity. Cell Metab 10, 260–272 (2009).

9. R. Karisch et al., Global proteomic assessment of the classical protein-tyrosine phosphatome and “Redoxome”. Cell 146, 826–840 (2011).

10. B. Boivin et al., Receptor Protein-tyrosine Phosphatase α Regulates Focal Adhesion Kinase Phosphorylation and ErbB2 Oncoprotein-mediated Mammary Epithelial Cell Motility. J Biol Chem 288, 36926–36935 (2013).

11. D. I. Brown, K. K. Griendling, Regulation of Signal Transduction by Reactive Oxygen Species in the Cardiovascular System. Circ Res 116, 531–549 (2015).

12. A. Mullard, FDA approves 100th small-molecule kinase inhibitor. Nat Rev Drug Discov 24, 891–895 (2025).

13. J. N. Andersen et al., Structural and evolutionary relationships among protein tyrosine phosphatase domains. Mol Cell Biol 21, 7117–7136 (2001).

14. Z. Y. Zhang, J. E. Dixon, Active site labeling of the Yersinia protein tyrosine phosphatase: the determination of the pKa of the active site cysteine and the function of the conserved histidine 402. Biochemistry 32, 9340–9345 (1993).

15. A. Salmeen et al., Redox regulation of protein tyrosine phosphatase 1B involves a sulphenyl-amide intermediate. Nature 423, 769–773 (2003).

16. R. L. van Montfort, M. Congreve, D. Tisi, R. Carr, H. Jhoti, Oxidation state of the active-site cysteine in protein tyrosine phosphatase 1B. Nature 423, 773–777 (2003).

17. C. L. Welsh, L. K. Madan, Protein Tyrosine Phosphatase regulation by Reactive Oxygen Species. Adv Cancer Res 162, 45–74 (2024).

18. A. D. Londhe et al., Regulation of PTP1B activation through disruption of redox-complex formation. Nat Chem Biol 16, 122–125 (2020).

19. L. V. Ravichandran, H. Chen, Y. Li, M. J. Quon, Phosphorylation of PTP1B at Ser(50) by Akt impairs its ability to dephosphorylate the insulin receptor. Mol Endocrinol 15, 1768– 1780 (2001).

20. A. D. Londhe, S. H. M. Rizvi, B. Boivin, In Vitro Activity Assays to Quantitatively Assess the Endogenous Reversible Oxidation State of Protein Tyrosine Phosphatases in Cells. Curr Protoc Chem Biol. 12, e84 (2020).

21. A. D. Londhe, B. Boivin, Measuring the Reversible Oxidation of Protein Tyrosine Phosphatases Using a Modified Cysteinyl-Labeling Assay. Methods Mol Biol 2743, 223– 237 (2024).

22. D. I. Brown, K. K. Griendling, Nox proteins in signal transduction. Free Radic Biol Med 47, 1239–1253 (2009).

23. K. M. Holmström, T. Finkel, Cellular mechanisms and physiological consequences of redox-dependent signalling. Nat Rev Mol Cell Biol 15, 411–421 (2014).

24. K. L. Milarski et al., Sequence specificity in recognition of the epidermal growth factor receptor by protein tyrosine phosphatase 1B. J Biol Chem 268, 23634–23639 (1993).

25. J. S. Biscardi et al., c-Src-mediated phosphorylation of the epidermal growth factor receptor on Tyr845 and Tyr1101 is associated with modulation of receptor function. J Biol Chem 274, 8335–8343 (1999).

26. Y. W. Lou et al., Redox regulation of the protein tyrosine phosphatase PTP1B in cancer cells. FEBS J 275, 69–88 (2008).

27. V. Vasquez-Montes, J. Gerhart, D. Thévenin, A. S. Ladokhin, Divalent Cations and Lipid Composition Modulate Membrane Insertion and Cancer-Targeting Action of pHLIP. J. Mol. Biol. 431, 5004–5018 (2019).

28. V. Vasquez-Montes et al., Ca2+-dependent interactions between lipids and the tumor-targeting peptide pHLIP. Protein Sci. 31, e4385 (2022).

29. B. Boivin, N. K. Tonks, Analysis of the redox regulation of protein tyrosine phosphatase superfamily members utilizing a cysteinyl-labeling assay. Methods Enzymol 474, 35–50 (2010).

30. H. Sies, Oxidative eustress: On constant alert for redox homeostasis. Redox Biol 41, 101867 (2021).

31. H. Zhou et al., The biological buffer bicarbonate/CO2 potentiates H2O2-mediated inactivation of protein tyrosine phosphatases. Chem Soc 133, 15803–15805 (2011).

32. M. Dagnell et al., Bicarbonate is essential for protein-tyrosine phosphatase 1B (PTP1B) oxidation and cellular signaling through EGF-triggered phosphorylation cascades. J Biol Chem 16, 12330–12338 (2019).

33. J. S. Lazo, K. E. McQueeney, J. C. Burnett, P. Wipf, E. R. Sharlow, Small molecule targeting of PTPs in cancer. Int J Biochem Cell Biol 96, 171–181 (2018).

34. A. Haque, J. N. Andersen, A. Salmeen, D. Barford, N. K. Tonks, Conformation-sensing antibodies stabilize the oxidized form of PTP1B and inhibit its phosphatase activity. Cell 147, 185–198 (2011).

35. J. V. Frangioni, P. H. Beahm, V. Shifrin, C. A. Jost, B. G. Neel, The nontransmembrane tyrosine phosphatase PTP-1B localizes to the endoplasmic reticulum via its 35 amino acid C-terminal sequence. Cell 68, 545–560 (1992).

36. F. G. Haj, P. J. Verveer, A. Squire, B. G. Neel, P. I. Bastiaens, Imaging sites of receptor dephosphorylation by PTP1B on the surface of the endoplasmic reticulum. Science 295, 1708–1711 (2002).

37. E. R. Eden, I. J. White, A. Tsapara, C. E. Futter, Membrane contacts between endosomes and ER provide sites for PTP1B-epidermal growth factor receptor interaction. Nat Cell Biol 12, 267–272 (2010).

38. I. A. Yudushkin et al., Live-cell imaging of enzyme-substrate interaction reveals spatial regulation of PTP1B. Science 315, 115–119 (2007).

39. Y. Romsicki, M. Reece, J. Y. Gauthier, E. Asante-Appiah, B. P. Kennedy, Protein tyrosine phosphatase-1B dephosphorylation of the insulin receptor occurs in a perinuclear endosome compartment in human embryonic kidney 293 cells. J Biol Chem 279, 12868– 12875 (2004).

40. V. Sangwan et al., Regulation of the Met receptor-tyrosine kinase by the protein-tyrosine phosphatase 1B and T-cell phosphatase. J. Biol. Chem. 283, 34374–34383 (2008).

41. K. Chen, M. T. Kirber, H. Xiao, Y. Yang, J. F. Keaney, Jr, Regulation of ROS signal transduction by NADPH oxidase 4 localization. J. Cell. Biol. 181, 1129–1139 (2008).

42. M. Geiszt, J. B. Kopp, P. Varnai, T. L. Leto, Identification of renox, an NAD(P)H oxidase in kidney. Proc. Natl. Acad. Sci. USA 97, 8010–8014 (2000).

43. A. Shiose et al., A novel superoxide-producing NAD(P)H oxidase in kidney. J Biol Chem 276, 1417–1423 (2001).

44. M. Menshikov et al., Urokinase plasminogen activator stimulates vascular smooth muscle cell proliferation via redox-dependent pathways. Arterioscler Thromb Vasc Biol 26, 801– 807 (2006).

45. S. S. Brar et al., An NAD(P)H oxidase regulates growth and transcription in melanoma cells. Am J Physiol Cell Physiol 282, 1212–1224 (2002).

46. E. C. Vaquero, M. Edderkaoui, S. J. Pandol, I. Gukovsky, A. S. Gukovskaya, Reactive oxygen species produced by NAD(P)H oxidase inhibit apoptosis in pancreatic cancer cells. J Biol Chem 279, 34643–34654 (2004).

47. L. Wang et al., Therapeutic peptides: current applications and future directions. Signal Transduct Target Ther 7 (2022).

48. M. Muttenthaler, G. F. King, D. J. Adams, P. F. Alewood, Trends in peptide drug discovery. Nat. Rev. Drug Discov 20, 309–325 (2021).

49. K. Imai, A. Takaoka, Comparing antibody and small-molecule therapies for cancer. Nat Rev Cancer 6, 714–727 (2006).

50. M. C. Smith, J. E. Gestwicki, Features of protein-protein interactions that translate into potent inhibitors: topology, surface area and affinity. Expert Rev Mol Med 14, e16 (2012).

51. D. Zane, P. L. Feldman, T. Sawyer, Z. Sobol, J. Hawes, Development and Regulatory Challenges for Peptide Therapeutics. Int J Toxicol 40, 108–124 (2021).

52. D. Thévenin, M. An, D. M. Engelman, pHLIPmediated translocation of membrane-impermeable molecules into cells. Chem Biol 16, 754–762 (2009).

53. M. An, D. Wijesinghe, O. A. Andreev, Y. K. Reshetnyak, D. M. Engelman, pH-(low)-insertion-peptide (pHLIP) translocation of membrane impermeable phalloidin toxin inhibits cancer cell proliferation. Proc. Natl. Acad. Sci. U.S.A. 107, 20246–20250 (2010).

54. K. E. Burns, M. K. Robinson, D. Thévenin, Inhibition of cancer cell proliferation and breast tumor targeting of pHLIP-monomethyl auristatin E conjugates. Mol Pharmaceutics 12, 1250–1258 (2015).

55. Y. K. Reshetnyak, O. A. Andreev, U. Lehnert, D. M. Engelman, Translocation of molecules into cells by pH-dependent insertion of a transmembrane helix. Proc. Natl. Acad. Sci. U.S.A. 103, 6460–6465 (2006).

56. D. Hanahan, R. A. Weinberg, The hallmarks of cancer. Cell 100, 57–70 (2000).

57. E. Bloch et al., Disrupting the transmembrane domain-mediated oligomerization of protein tyrosine phosphatase receptor J inhibits EGFR-driven cancer cell phenotypes. J. Biol. Chem. 294, 18796–18806 (2019).

58. J. Gerhart, A. F. Thévenin, E. Bloch, K. E. King, D. Thévenin, Inhibiting Epidermal Growth Factor Receptor Dimerization and Signaling Through Targeted Delivery of a Juxtamembrane Domain Peptide Mimic. ACS Chem. Biol. 13, 2623–2632 (2018).

59. J. Wehr et al., pH-Dependent Grafting of Cancer Cells with Antigenic Epitopes Promotes Selective Antibody-Mediated Cytotoxicity. J. Med. Chem. 63, 3713–3722 (2020).

